# Enteric Viruses and Free-Living Amoebae: Protozoa as Potential Reservoirs and Transport Vessels for human Norovirus and Adenovirus

**DOI:** 10.1101/2025.04.07.647535

**Authors:** Mats Leifels, Rafik Dey, Alyssa Wiedmeyer, Cheng Dan, Claudia Kolm, Fuqing Wu, Kwanrawee Sirikanchana, Andreas H. Farnleitner, Nicholas J. Ashbolt

## Abstract

Human norovirus (HNoV) and human adenovirus (HAdV) are major causes of acute viral gastroenteritis globally and environmentally transmitted via the faecal-oral route through contaminated food and water. Recent evidence of these enteric viruses residing within environmental free-living amoebae (FLA)—specifically Vermamoeba vermiformis, Acanthamoeba polyphaga, and Willaertia magna—has significant implications for environmental virology and public health. The incorporation of HNoV into the cytoplasm and vacuoles of V. vermiformis and A. polyphaga, as well as the nuclear localization of HAdV within W. magna, was demonstrated using quantitative PCR and fluorescence microscopy. Intact HNoV and HAdV virions persisted inside FLA trophozoites, cysts, and extracellular vesicles for up to 12 days. Moreover, HAdV retained infectivity in buffalo green monkey kidney cells following intracellular persistence, suggesting these viruses can evade amoebal digestion and structural degradation. In the case of HAdV, nuclear incorporation, preservation of capsid integrity, and detection of mRNA associated with adenoviral fiber protein synthesis further suggest the possible initiation of virus-related transcriptional activity within the amoeba host.

These findings challenge current assumptions about virus removal rates in sewage treatment, food safety protocols, and drinking water production. The enhanced persistence and protection conferred by FLA may also impact microbial risk assessments for recreational water use, particularly in sewage-impacted rivers and lakes. Recognition of FLA as reservoirs and transport vessels for enteric viruses necessitates a re-evaluation of existing water and sanitation guidelines to better mitigate environmental transmission pathways.

**Graphical Abstract:** 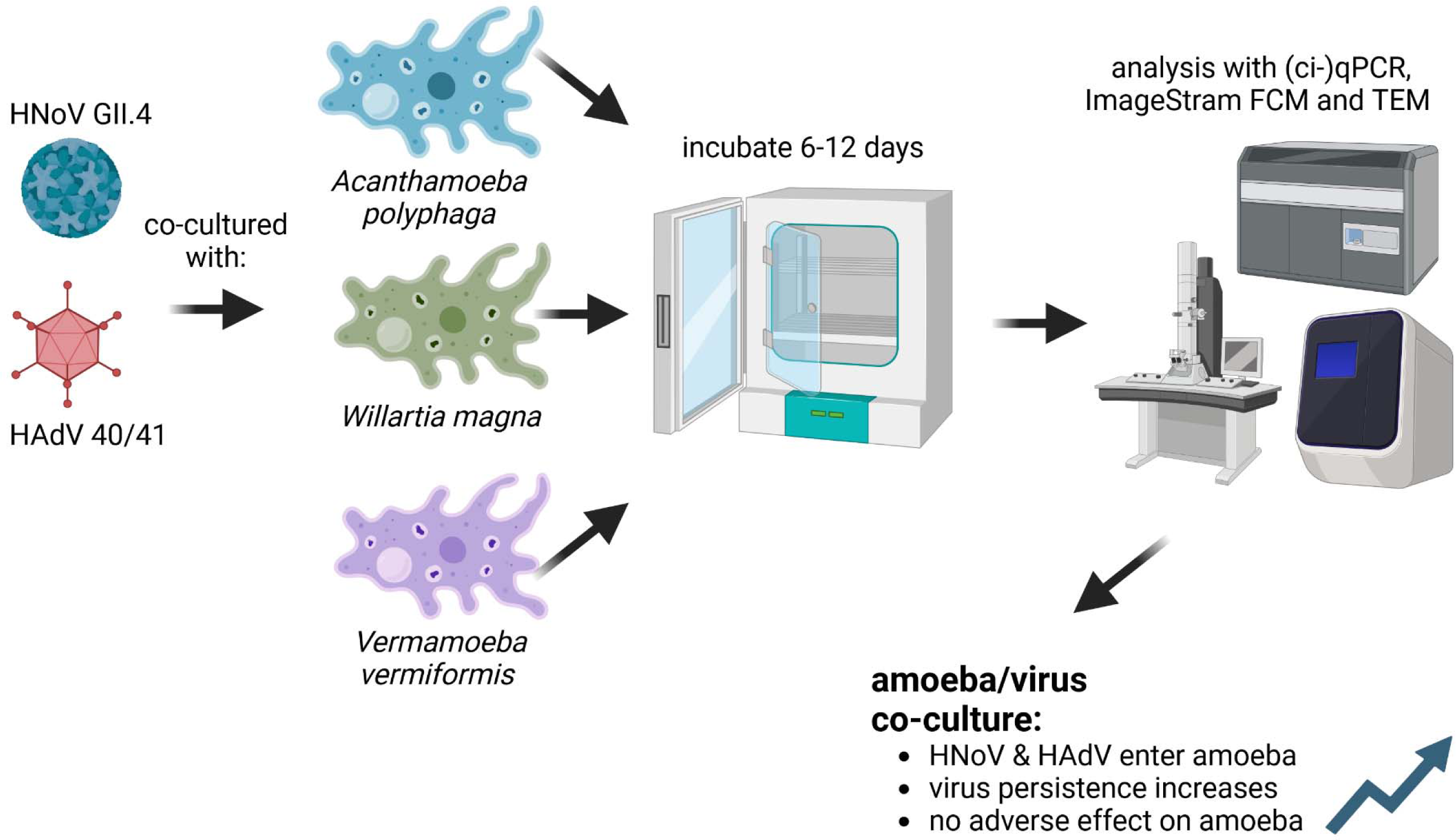

## Introduction

Acute and chronic illnesses associated with food and water are widespread and contribute to an enormous burden of disease and economic strain in developing and developed countries (Cassini et al., 2016; Chen et al., 2023; Ko et al., 2005). According to the United States Centers for Disease Control and Prevention (US-CDC), 31 major pathogens, spanning bacteria, viruses and parasitic protozoa were responsible for approximately 9.9 million annual cases of food- and waterborne infections in 2019 and the US alone (Scallan Walter et al., 2025). While the number of disability adjusted life years and overall mortality of diarrheal disease resulting from unsafe water decreased from 1990 to 2019 by 50% and 59%, respectively, it is still among a leading cause of death across all age groups (Chen et al., 2023; Troeger et al., 2018).

Among microbial waterborne pathogens associated with diarrhoea, human norovirus (HNoV) is the most common and causes at least 500-800 yearly deaths, 71,000 hospitalizations and 1.9 million outpatient clinic visits in the US, as well as leading to more than 2 billion USD of economic loss due to healthcare expense and lost productivity (Bányai et al., 2018). In addition to the absence of an available vaccine, the high infectivity and low infectious dose of human norovirus (HNoV) contribute to its widespread transmission. As few as 10 viral particles (Hutson et al., 2004) or 5 genome copies (Atmar et al., 2008) may cause infection, while infected individuals shed between 9 and 11 log₁₀ genome copies per gram of stool—often irrespective of symptom onset (Free et al., 2019; Lee et al., 2007).

Human adenoviruses (HAdV) are another group of enteric viruses capable of causing acute gastrointestinal infections (GI), particularly following exposure to faecally contaminated water—for example, during recreational activities (Girardi et al., 2019; Leifels et al., 2019). While not as infectious as HNoV, and more often resulting in asymptomatic or mild GI and respiratory infections, worldwide, GI-associated HAdV subtypes HAdV40 and HAdV41 have been reported to cause 5-20% of acute gastroenteritis among children under the age of 5 years (Ziros et al., 2015) with the highest prevalence at 20% in Africa (Khales et al., 2024).

Once excreted by the human host, both HNoV and HAdV exhibit high environmental persistence, remaining stable under conditions such as sunlight exposure, temperature fluctuations, and pH changes, while retaining infectivity for several days to weeks (Hall, 2012; O’Shea et al., 2019). For example, HAdV infectivity has been shown to remain stable in both surface and drinking water for up to 70 days at 4°C and 20°C (Strathmann et al., 2016), highlighting its potential to pose a significant health risk in untreated water associated with recreational activities. Similarly, the prolonged persistence of HNoV on surfaces and fomites contribute to its frequent association with healthcare-related outbreaks (Lei et al., 2017) .

Free-living Amoebae (FLA) are unicellular eukaryotic protozoa that predominantly feed on bacteria within water and soil environments. Due to their broad ecological adaptability, FLA can thrive in both natural and engineered water systems, including riverbeds, tidal zones, cooling towers, hospital water networks, water treatment plants, and premise plumbing systems (Ashbolt, 2023; Samba-Louaka et al., 2019; Thomas and Ashbolt, 2011) as well as throughout wastewater treatment facilities worldwide (da Silva et al., 2024). Depending on environmental conditions such as stress levels and nutrient availability, FLA alternate between two life stages: an actively growing trophozoite form, in which they feed primarily through phagocytosis via pseudopod extension and retraction, and a dormant cyst form, which allows for long-term survival under adverse conditions. In their dormant cyst form, FLA have been demonstrated to be most resistant against microbicidal UV and various chemical disinfection treatments such as hypochlorite and monochloramine, all commonly used for wastewater and drinking water treatment (Anwar et al., 2018). The remarkable resilience of FLA is exemplified by the successful revival of cysts from the Flamella genus that had been preserved in permafrost for up to 35,000 years (Shmakova et al., 2016) and Acanthamoeba cysts remaining viable after treatment with 100 mg/l chlorine (free and combined) for 10 min, as well as 80 °C thermal treatment (Storey et al., 2004).

Amoeba-resisting microorganisms (Kebbi-Beghdadi and Greub, 2014) also persist within the protection of FLA trophozoites, extracellular vesicles and cysts (Folkins et al., 2020b); (Shaheen and Ashbolt, 2018), many of which have become accidental human pathogens, particularly for susceptible individuals in hospitals and care facilities (Buse et al., 2017; Collier et al., 2021; Hamilton et al., 2016). As opportunistic pathogens, FLA themselves are of public health importance due to the wide range of associated diseases such as keratitis or amebiasis and their omnipresence in water networks that leads to direct contact with susceptible individuals (Muchesa et al., 2016). Interactions between FLA and opportunistic/obligate pathogenic bacteria such as Legionella pneumophila are well established (Flemming and Wingender, 2010; Flemming and Wuertz, 2019). The relatively recent discovery of a close relationship between FLA and (giant) viruses (Scheid, 2015) lead to an increasing interest in the co-existence between viruses and amoeba (Hsueh and Gibson, 2015; Verani et al., 2016; Zhang et al., 2022).

First insights into enteric virus-FLA interaction were gained for highly abundant Acanthamoeba spp., which could have been shown to contain HAdV, potentially due to the structural molecular similarities between surface proteins in the amoeba nucleus membrane and receptors present on the surface of human macrophages (Sun et al., 2018). Similar behaviour could be observed for HAdV within Willaertia magna, another abundant FLA (Chaúque and Rott, 2022; Fan et al., 2024). Once they entered their amoeba carrier, adenoviruses were shown to utilize them as vehicle and reservoir, thus increasing their persistence towards environmental influences while not decreasing their infectivity (Scheid and Schwarzenberger, 2012). Acanthamoeba spp. has also been shown to incorporate particles of the HNoV surrogate murine norovirus (MNV) under laboratory conditions, which led to a notable increase in their already high resistance towards chemical disinfectants (Hsueh and Gibson, 2015). Preliminary studies indicate the possibility that HAdV might even be able to replicate inside the nucleus, which could potentially have implications for quantitative microbial risk assessment of enteric viruses in general (Colson et al., 2017; Mattana et al., 2006).

Atanasova et al. (2018) were able to show coxsackievirus B5 entering FLA of the species V. vermiformis and A. polyphaga (both isolated from hospital water taps). The virions avoided consumption by the FLA and were found accumulating in vacuoles before their excretion by the amoeba host via vesicles. While the interactions between FLA and various enteric and environmental viruses have received growing attention over the past decade, a direct association between FLA and human norovirus (HNoV) is yet to be described. Given the global public health significance of HNoV and the frequent co-occurrence of high concentrations of both organisms in sewage-impacted waters—particularly in secondary wastewater treatment plants and sewage-contaminated rivers and lakes (da Silva et al., 2024; Walker et al., 2024) – FLA-HNoV interactions seem likely, if not also potential autochthonous replication of HNoV within FLA of freshwater and marine environments.

If FLA-human virus interactions occur to a significant degree, that would necessitate a reassessment of pathogen removal targets in sewage treatment facilities to ensure minimal risk to downstream receiving waters used for recreational activities or drinking water production. Given the well-documented limitations of traditional faecal indicator bacteria such as Escherichia coli and enterococci, HAdV—along with bacteriophages like crAssphage—have been proposed as more reliable viral indicators of water quality (Gómez-Gómez et al., 2024) However, if differential behaviours occur through FLA interactions, that could potentially change their suitability for that role. Herein we investigated the effects of co-cultured HAdV and HNoV with FLA to assess their environmental persistence.

## Materials and Methods

### Amoeba Cultures and Maintenance

FLA Vermamoeba vermiformis (ATCC 50237), Acanthamoeba polyphaga (ATCC 30461) and Willartia magna (ATCC 50035) were acquired from the American Type Culture Collection. Amoeba were cultured in 5cm^3^ non-vented flasks with 5mL of Serum Casein Glucose Yeast Extract Medium (SCGYEM; ATCC 1021), according to the manufacturer’s recommendations. The medium was exchanged every 72 – 96 hours to maintain active trophozoites in culture. Amoebae were gently detached from the flask by drawing up the newly added medium and running it three times over the bottom of the flask and maintained at 25 °C in the dark. Before each virus-amoeba co-culture, a new subculture was prepared one to two days before intended use to ensure actively growing trophozoites were used.

### Adenovirus cell-culture

Human adenovirus HAdV40/41 strains and Buffalo Green Monkey (BGM) cells were kindly provided by Dr. Xiaoli Lilly Pang, Provincial Laboratory for Public Health in Edmonton, AB, Canada. HAdV40/41 was cultured as described in our previous work (Leifels et al., 2016). In brief, BGM cells were grown in ventilated culture flasks using Dulbecco’s MEM (Sigma-Aldrich Co., MO, USA) supplemented with 10% heat inactivated fetal bovine serum (FBS) and 1% penicillin-streptomycin to inhibit bacteria growth.

### FLA - virus co-cultures

HNoV GII.4 aliquots were kindly provided by Dr. Xiaoli Lilly Pang, Provincial Laboratory for Public Health in Edmonton, AB, Canada. To prepare the FLA for virus co-culture, amoebae were gently detached from the flask by drawing up the newly added medium and running it three times over the bottom of the flask. Media was transferred to 15mL conical tubes then cells were counted under the microscope on a haemacytometer (Merck-Millipore, ON, Canada). HNoV originating from stool and HAdV (both infectious and inactivated by incubation at 95° C for 10 Min) were added at a concentration of approximately 1×10^7^ virions/mL to amoebae at 10^5^ amoebae/mL to reach a MOI of 100. Co-culture tubes were then incubated at 37 °C in the dark, and aliquots were extracted at days 0, 1, 2, 3, and 6 for HNoV and 0, 1, 2, 3, 4 and 12 for HAdV. Trophozoites and cysts were again counted from duplicates using a haemacytometer. Tubes were then transferred to −20°C until further processing.

### Integrated Cell-Culture (ICC)-qPCR

For the ICC-qPCR, cells were transferred from the flask to 6-well tissue culture plates. After 48 hours at 37 °C and in the presence of 5% CO_2_, the medium containing FBS was removed, the wells were washed with sterile filtered PBS and replicate plates were infected with 300 µL of HAdV in medium without FBS. After inoculation for 90 minutes, half of the plates were transferred to −20 °C while the remaining plates were supplemented with maintenance medium containing 2.5% FBS. After an incubation period between 96 to 110 hours, the cells were transferred to −20 °C. BGM cells were detached from the plate surface by three cycles of freezing and thawing. After that, nucleic acids from the cell lysate were extracted and viral genome copies were determined by qPCR. An increase of the nucleic acid copy number between the 90 minutes and 110 hours plates by the factor 10 or more was considered as positive for infectious HAdV.

### PMA capsid-integrity qPCR (PMA-ci-qPCR)

After co-culture with W. magna, 200 µL aliquots of the washed and resuspended pellet were treated with PMA (Biotium Inc., CA, USA) using a final concentration of 0.04 mM as described in our previous study (Leifels et al., 2019). Tubes were mixed gently by inverting several times before incubation at room temperature for 30 min in the dark and 10 min in daylight before being exposed to 4-Watt Blue Light for 15 Min. Positive and negative controls as well as qPCR interreference by PMA were conducted as described before (Leifels et al., 2021; Leifels et al., 2016).

### Genome extraction and cDNA synthesis

Virions internalized by the FLA were extracted by lysing the amoebae using freeze thaw cycles and syringe pressure/rupture as described in our previous work (Atanasova et al., 2018). Freeze thaw cycles consisted of 25 °C and −80 °C, repeated twice. Samples were then passed continuously and repeatedly through a 20-gauge 3 mL needle (BD, ON, Canada). The whole genome DNA was extracted from aliquots of 200 µl using the QIamp Blood and Tissue Kit (Qiagen, ON, Canada) according to the manufacturers protocol. 10 µl of the eluate was immediately converted from RNA into cDNA using the High-Capacity cDNA Reverse Transcription Kit (Applied Biosystems, ON, Canada) and stored in 20°C until further analysis.

### Detection of HAdV messenger RNA

To aid in the differentiation between infectious and non-infectious HAdV, messenger RNA (mRNA) involved in the formation of the virus capsid fiber instead of genomic DNA was targeted, as proposed by Ko et al. (2005). The assumption being that mRNA target would only be present during HAdV40/41 replication inside their FLA host. The primer and probe sequences used are listed in Table 1 and the cycle conditions are as follows: 95°C for 3 min, followed by 50 cycles of a 95°C denaturation for 15 s, 55°C annealing for 5 s, and extension at 72°C for 10 s.

**Table 1:**
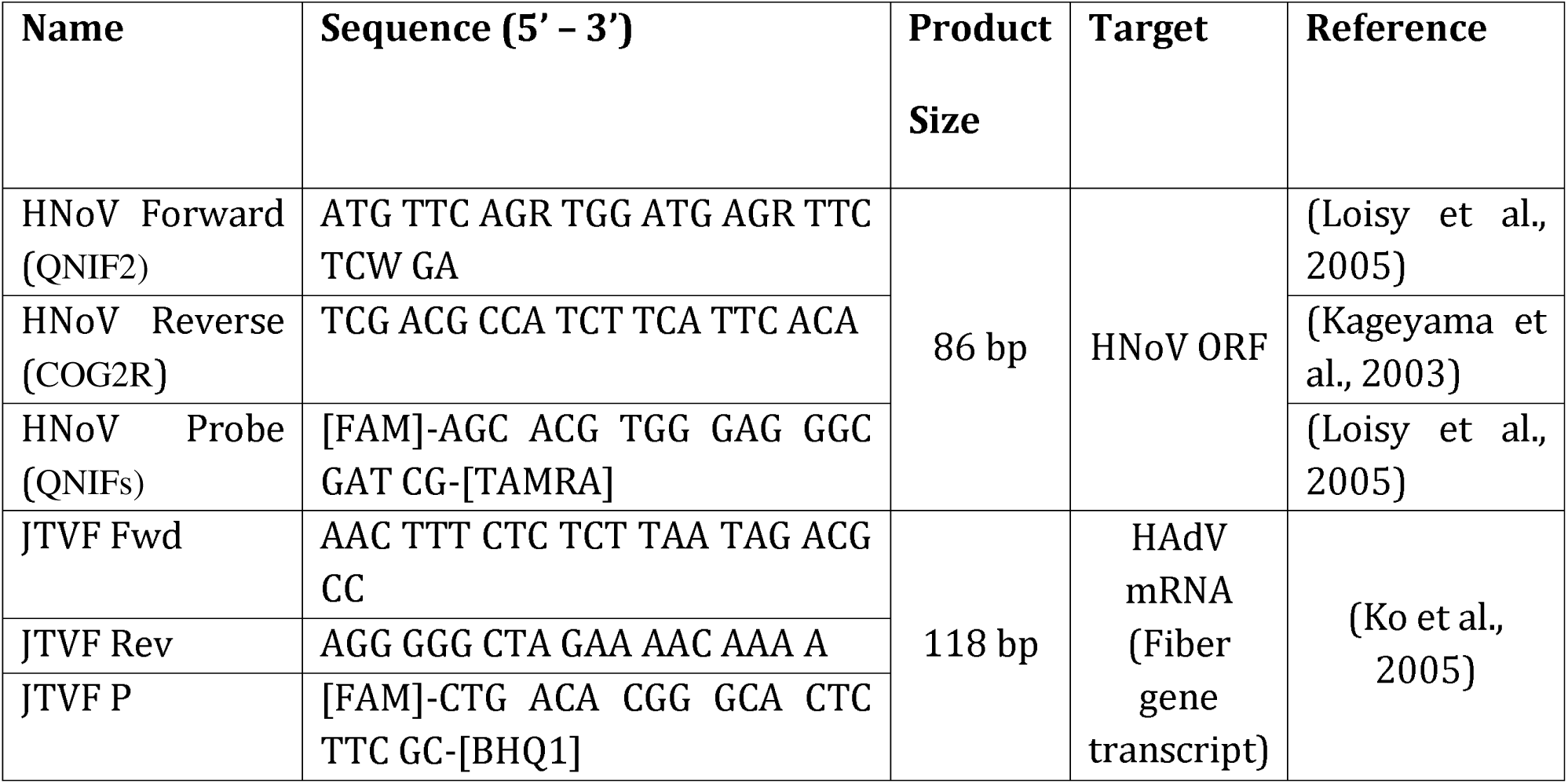
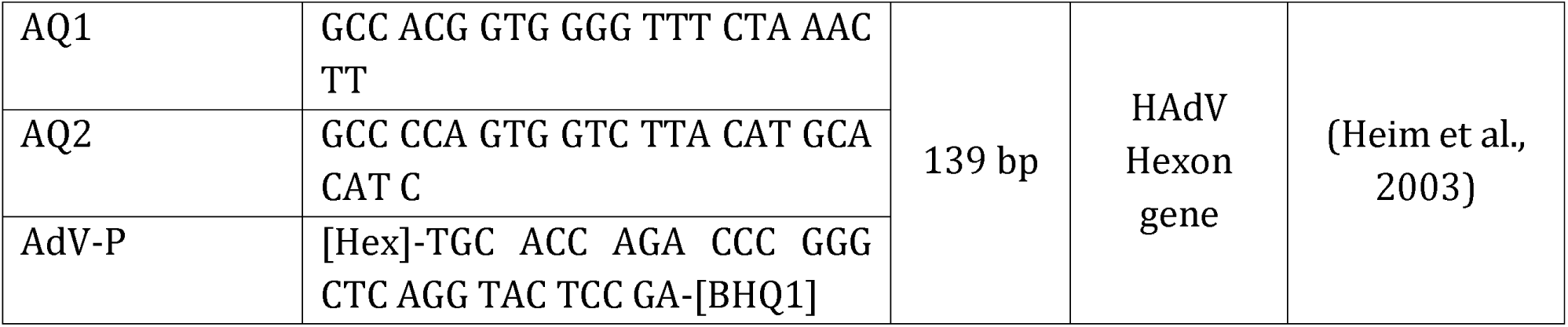
Sequences of Primers and probe used for HNoV GII.4, HAdV40/41 and HAdV40/41 mRNA specific qPCR.

### Quantitative PCR Analysis for HAdV and HNoV and FLA

The quantitative PCR (qPCR) was undertaken using the TaqPath qPCR Master Mix (Thermo Fisher, MA, USA) on a Qiagen Rotor-gene Q cycler (Qiagen, ON, Canada) following the protocol published by the International Standards Organization (ISO, 2017). Limit of detection (LOD) in copies per qPCR reaction was determined in serial dilutions of known genBlock standards (Integrated Genome Technologies, ON, Canada) and calculated to be 25 genomic copies for HAdV and 50 genomic copies for HNoV. Cycling conditions were as follows for HNoV: preheating for 3 min at 95°C followed by 45 cycles of 15 s at 95°C for denaturation and 1 Min at 60°C for annealing and 1 min at 65°C for extension. HAdV, 95 °C for 3 min, then 45 cycles of 95°C for 15 s, 60 °C for 1 min, and 78 °C for 5 s to acquire the fluorescence. Primer and probe sequences are listed in Table 1.

### Transmission Electron Microscopy (TEM)

To allow the visualization of virus incorporated within the three different FLA, co-cultures were prepared for the three species of amoebae with an increased MOI of 1,000 while amoebae without the addition of HNoV or HAdV were used as a control. After 48 hours of co-incubation, virus-amoeba mixtures were pelleted using low speed centrifugation, the supernatant was discarded, and the culture was resuspended in electron microscopy fixative (Electron Microscopy Sciences, PA, USA) before samples were submitted to the Advanced Microscopy Facility (University of Alberta AB, Canada) for transmission electron microscopy imaging.

### ImageStream flow cytometry

For analysis of the incorporation of HNoV GII.4 and HAdV40/41 into FLA, virus cultures were stained with SYBR^®^ Green II (for HNoV and HAdV) and Hoechst 33342 for DNA (for HAdV) (Thermo Fisher, MS, USA) for 24 hours before pelleting the mixture in a centrifuge at low speed. The pellet was then washed three times with sterile filtered PBS using Amicon 100K Centrifugal filter units (Merck-Millipore, ON, Canada) to remove excess stain. Aliquots of the stained virions were then used for co-culture with FLA in 15 mL conical tubes, as previously described (Folkins et al., 2020b). After 12 h, amoebae were transferred from the culture flask into separate 1.5 mL reaction tubes and analyzed using an the Amnis

ImageStream Mark II (Merck-Millipore, ON, Canada) at 494 nm. FLA cultured in the absence of viruses were used as a negative control and to determine autofluorescence.

### Statistical analysis

Statistical analysis and visualization of all experiments was performed in Microsoft Excel (Microsoft Inc., CA, USA) and GraphPad Prism (GraphPad Software, CA, USA). P-values below the cutoff of 0.05 were determined to be statistically significant.

## Results

### Potential adverse effects of HNoV on FLA growth rates

Co-cultures with HNoV (all three FLA species) and HAdV (only with Willaertia magna) were undertaken to determine if the presence of viruses could be associated with an adverse amoebae growth rate. However, no consistent difference for FLA trophozoites and cysts over time when compared to the virus-free amoebae controls (Figure 1A-C), inferring no apparent growth-impact with the internalization of enteric virions.

**Figure 1:**
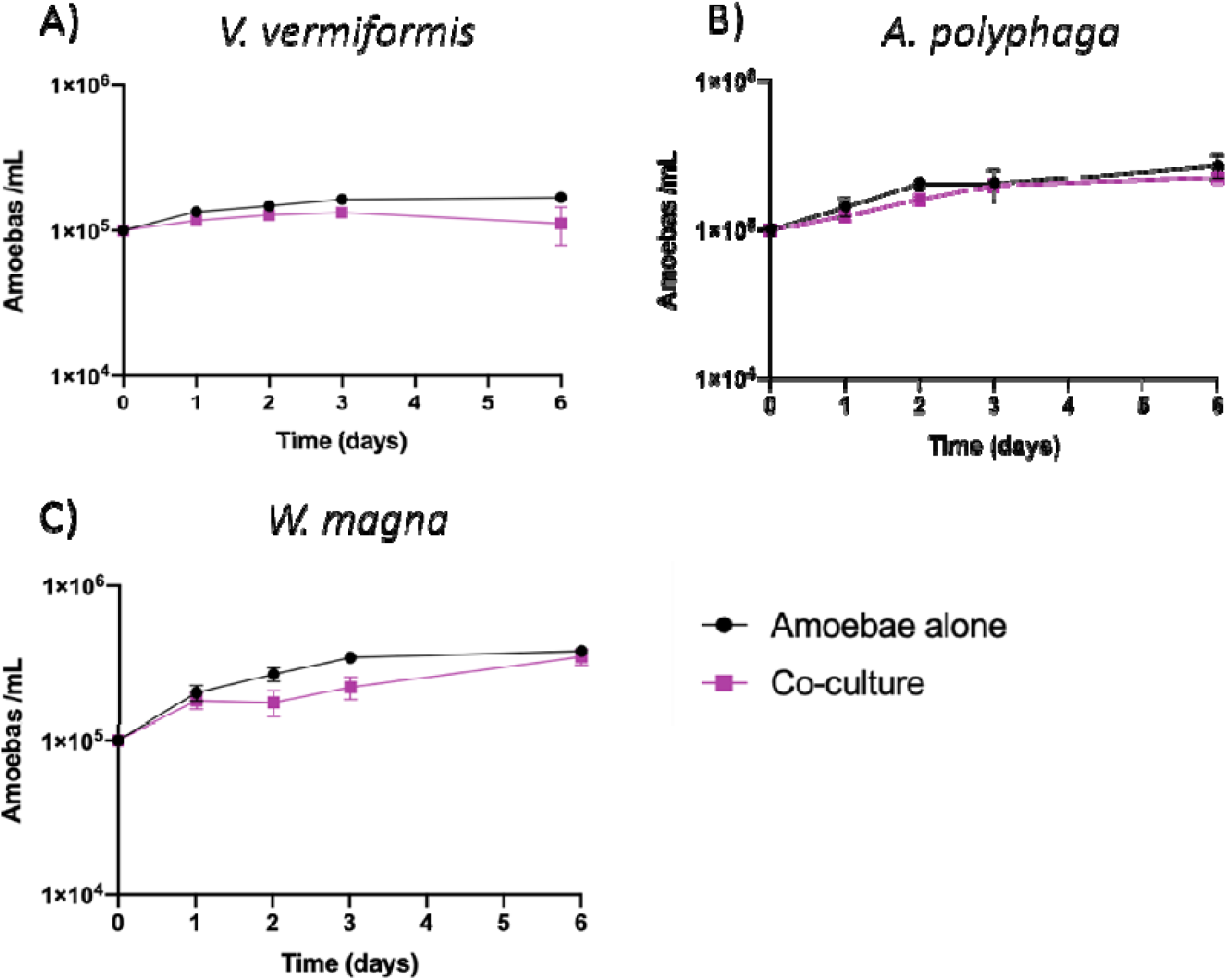
Growth rates of V. vermiformis (A), A. polyphaga (B), and W. magna (C). FLA and HNoV were co-cultured at a MOI of 100 and incubated at 37 °C in the dark. Aliquots were extracted daily until day 6, and trophozoites and cysts were quantified using a hemocytometer. Controls consist of amoebae without exposure to viruses. Error bars indicate standard deviation (n=4).

### HNoV persistence in FLA

Haemocytometer quantification of all three FLA in the presence of HNoV further indicated no negative effect by co-culture (Figure 2). All co-cultures amoebae under investigation showed no notable decrease over the six-day period when compared to virus-free control that were conducted separately under identical conditions (data not shown).

**Figure 2:**
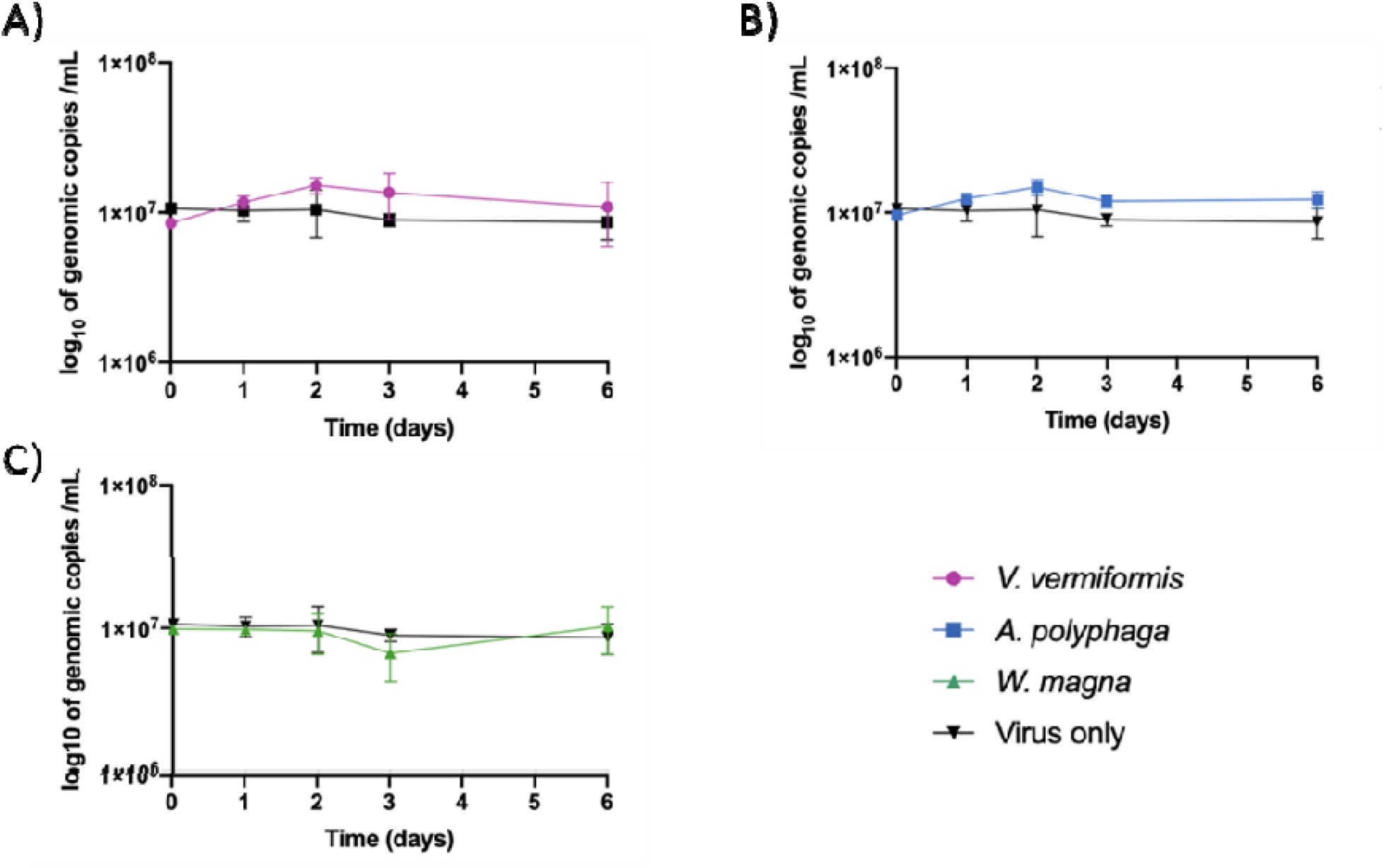
Genome concentrations of HNoV when co-cultured with V. vermiformis (A), A. polyphaga (B) and W. magna (C) and cultivated separately. FLA and HNoV were co-cultured at a MOI of 100 and incubated at 37 °C in the dark. Aliquots were extracted daily until day 6, and trophozoites and cysts were quantified via haemocytometer, virions via qPCR. Error bars indicate standard deviation (n=4).

### Transmission Electron Microscopy (TEM) analysis

Using TEM to visualise internalised HNoV and HAdV within 48 h co-cultures with FLA identified virion-like structures. (Figure 3) For HNoV, distinct accumulations of structures resembling virions can be seen in aggregates within the cytoplasm of V. vermiformis and A. polyphaga (see Figure 3 B and D) while no HNoV virus-like particles were observed in the nucleus or nuclear membrane of co-cultured FLA. The shape and structure of organelles in both FLA that internalized HNoV were indistinguishable from those in amoebae controls without the virus, further supporting that viron internalization produced no observable morphological (cell pathogenic) changes or cell injury. Higher magnification images of V. vermiformis and A. polyphaga showed virions within the amoebae cytoplasm (Figure 4). Individual particles appear to have icosahedral-shapes, and their measured size was comparable to that reported for intact HNoV (30-40 nm).

**Figure 3:**
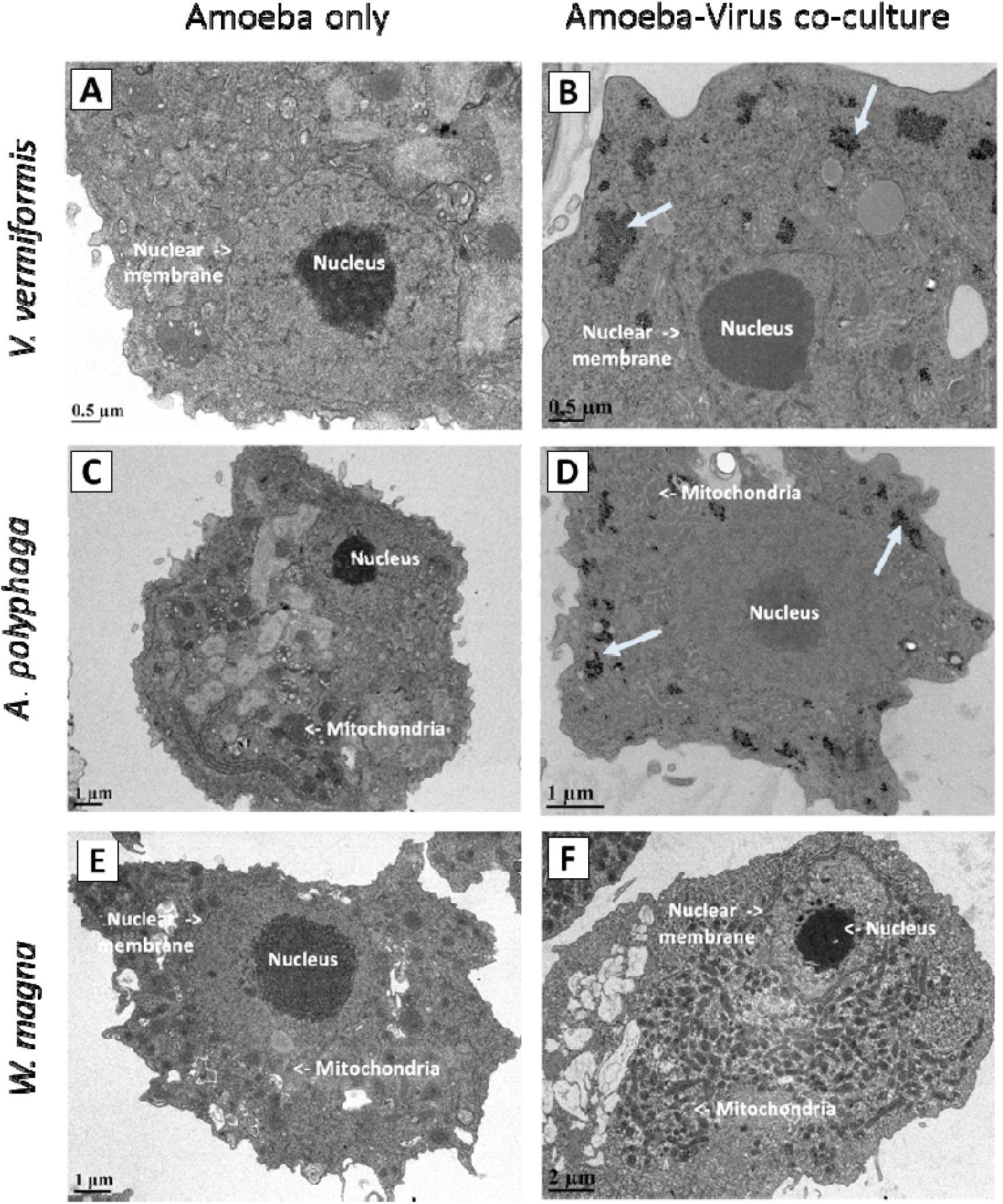
Transmission Electron microscopy of the FLA cultured with (A, C and E) and without (B, D and F) HNoV. After growing with and without HNoV for 48 hours, internal organelles such as the nucleus, nuclear membrane, and mitochondria are (indicated on individual images) show no cytopathic effect or adherent growth. Clusters of aggregated virus-like particles are indicated by the white arrows in V. vermiformis (B) and A. polyphaga (D) co-cultures, while n HNoV virions could be observed inside W. magna (F).

**Figure 4:**
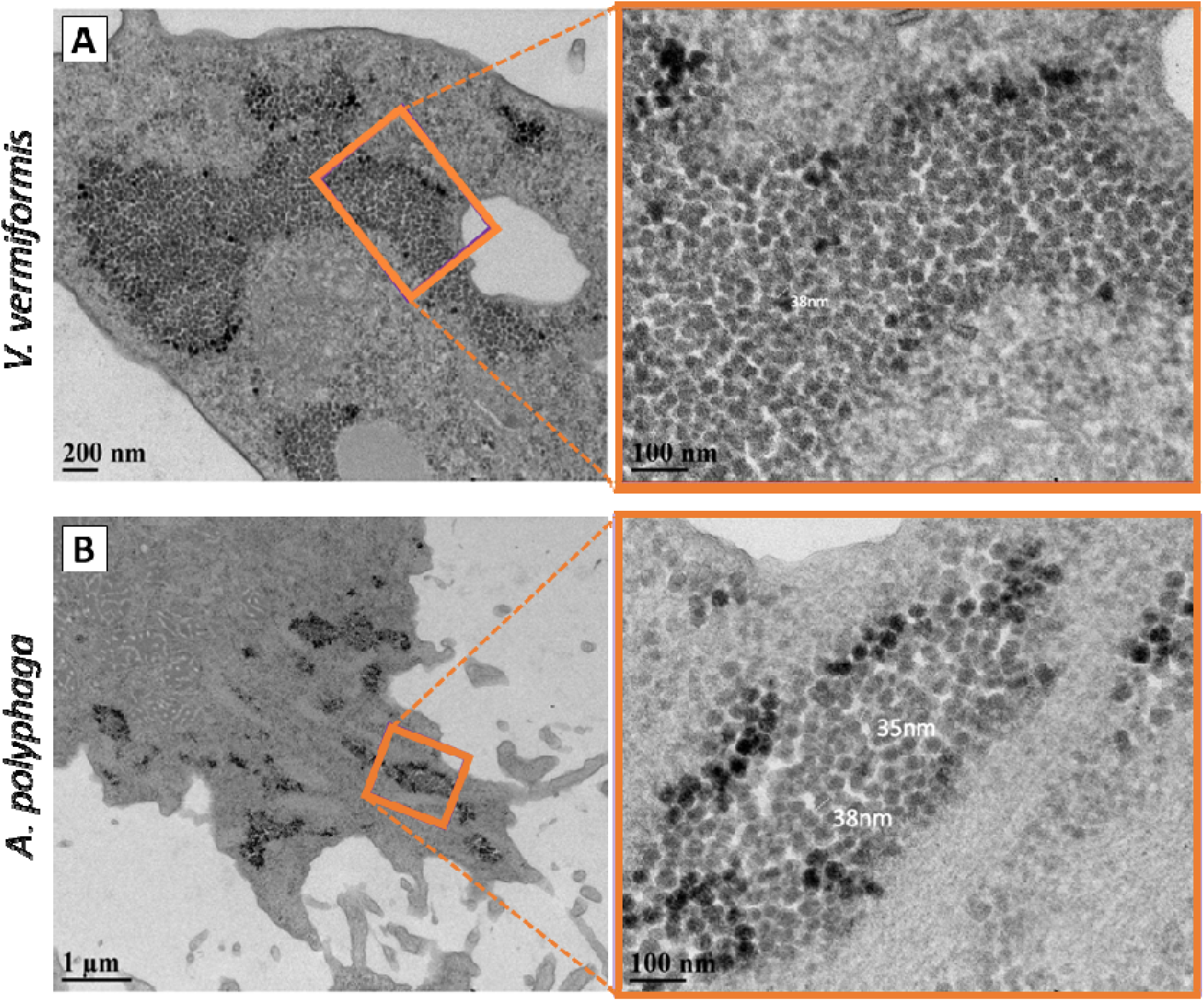
Magnified TEM images of V. vermiformis (A) and A. polyphaga (B) co-cultured with HNoV. After growing i the presence of presumably infectious virions for 48 hours, clusters of aggregated virus-like particles can be seen compartmentalized in the cytoplasm. Magnified images show structures that resemble individual virus-like particles in size and shape.

Similarly, clusters of icosahedrals that resembled HAdV in size and structure were present in W. magna after incubation of this FLA in the presence of infectious viruses fo 48 h (Figure 5). Unlike HNoV, though, HAdV virions appeared to have passed the nucleus membrane and resided within the amoeba’s nucleus. Comparable to HNoV, no phenotypical changes commonly associated with cytopathic damage were observed for the amoebae when cultured either with or without HAdV.

**Figure 5:**
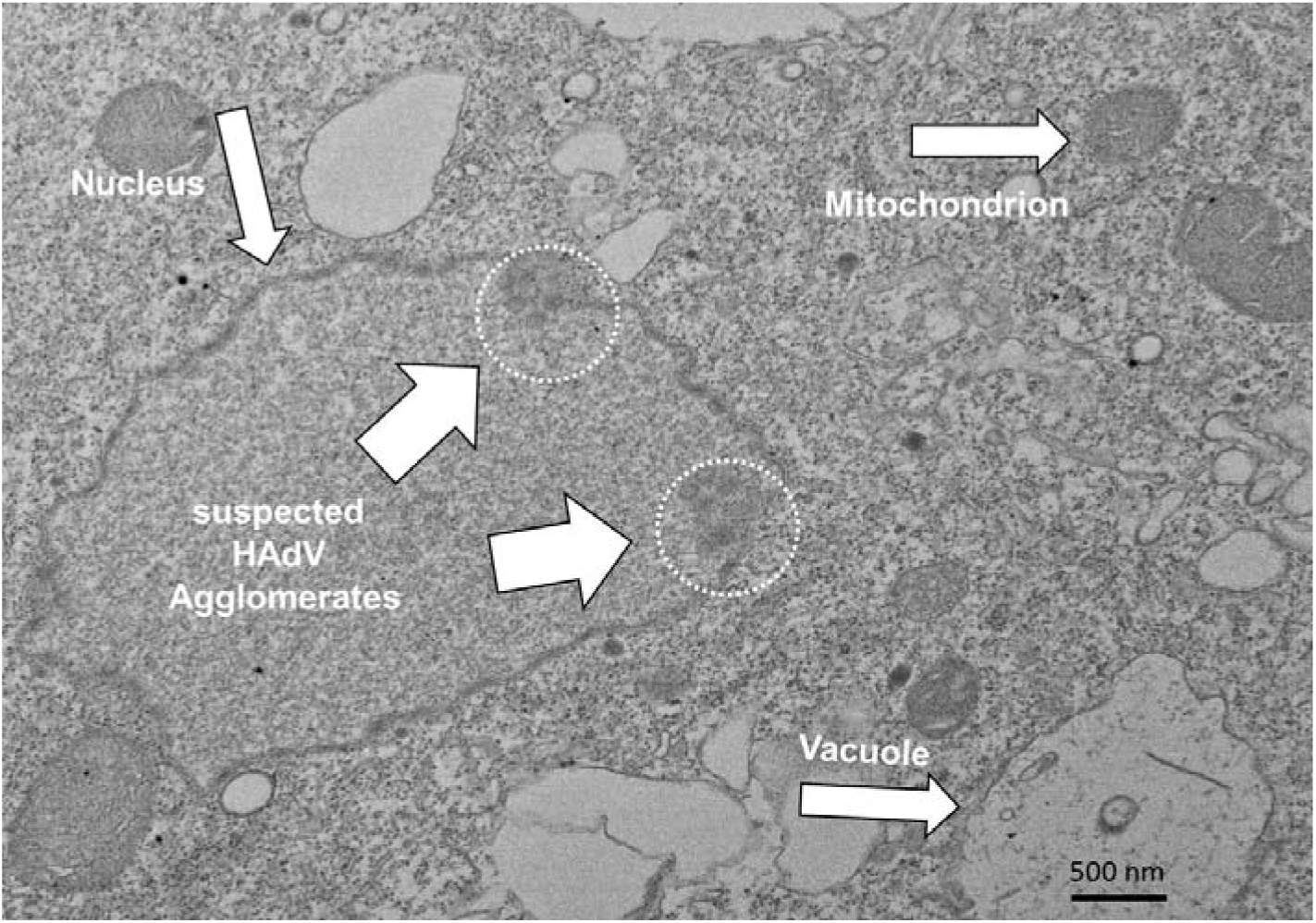
Transmission Electron microscopy of W. magna co-cultured with infectious HAdV. After growing wit HAdV for 48 hours, organelles such as the nucleus and mitochondria as well as cellular structures like the nuclear membrane and vacuoles show no cytopathic effect or adherent growth. Suspected agglomerates of aggregated HAdV virions are visible inside the amoeba nucleus, indicating the ability of HAdV to pass through this supposedly impermeabl barrier and open the possibility of virus propagation.

### ImageStrean Flow Cytometry

The ImageStream combines flow cytometry and confocal laser scanning microscopy and allowed for the localization of HNoV internalized within all three FLA as well as HAdV within W. magna. HNoV virions stained with SYBR^®^ Green II RNA stain were co-cultured with the respective FLA species for 12 hours before introduction into ImageStream. Simila to TEM results, both V. vermiformis and A. polyphaga images showed internalization of HNoV virions (see Figure 6). However, unlike TEM, staining with genome reactive dye indicates the presence of HNoV within the cytoplasm of tryphozoites and vesicles of W. magna two days after co-culture (see Figure 6C).

**Figure 6:**
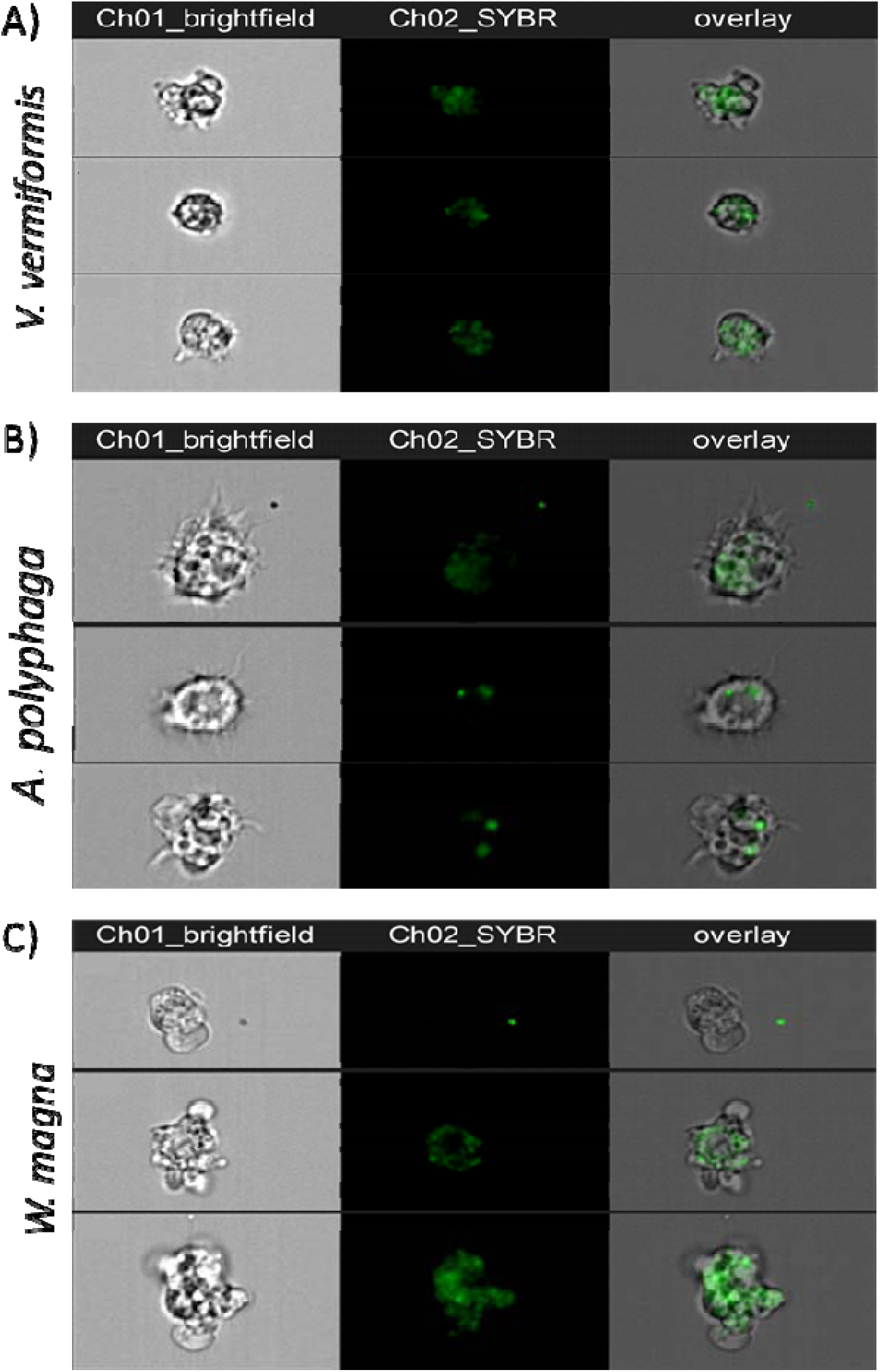
ImageStream of V. vermiformis (A), A. polyphaga (B), and W. magna (C), cultured with and without HNoV. Virions were stained with SYBR green II before inoculation with FLA. A minimum of 500 cells were captured per sample and analysis was performed using the Luminex IDEASTM suite.

To evaluate the presence of HAdV virions inside the amoeba nucleus, viruses were stained with the nucleus permeating Hoechst dye (blue), while both W. magna trophozoite and vesicles were stained with SYBR^®^ Green II (green). Overlap of both the green and blue spectra thus allowed to depict HAdV that entered the nucleus as well as those that were supposedly actively excreted by the amoebae in vesicles (see Figure 7).

**Figure 7:**
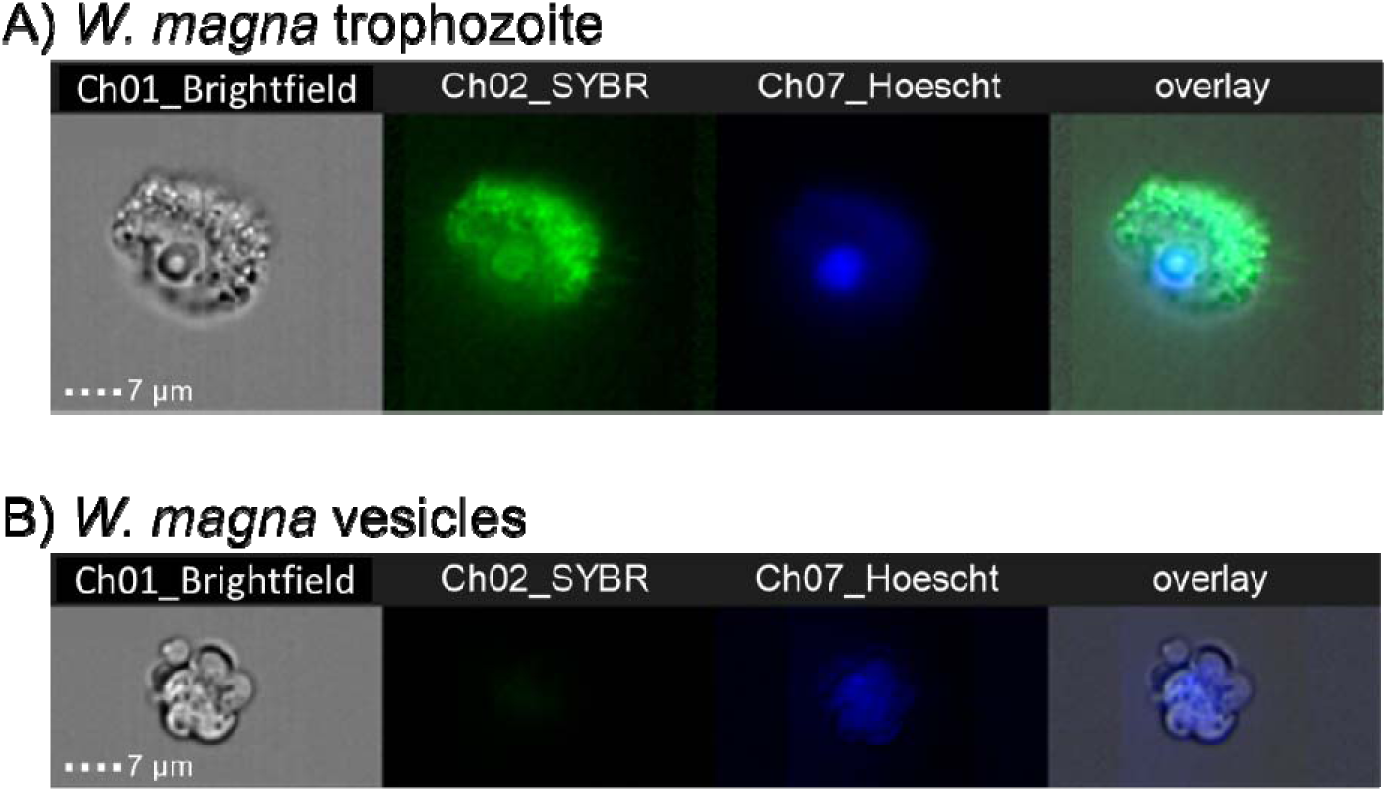
ImageStream of W. magna trophozoites (A) and vesicles (B) co-cultured with infectious HAdV. Befor inoculation, virions were stained with Hoechst and the amoeba culture with SYBR Green II for 24 hours to allow for the visualization of viruses that crossed the nuclear membrane and entered the supposedly impermeable amoeba nucleus.

### Capsid-integrity and mRNA qPCR to determine HAdV replication activity

After identification of HAdV incorporated into the nucleus of W. magna, the reproductive activity of the infectious and heat-inactivated virions was evaluated by targeting messenger RNA associated with the formation of HAdV fibers as well as the integrity of the adenoviral capsid. As shown in Figure 8, co-culture of W. magna with infectious HAdV virions showed no reduction in genomic copies assayed using ICC-qPCR and PMA-ci-qPCR for the 12-day experiments. Furthermore, mRNA – while under the limit of quantification - was detectable in all samples. When HAdV was heat-inactivated before co-culture with the FLA, a reduction of genomic copies was observed starting immediately after inoculating co-cultures.

**Figure 8:**
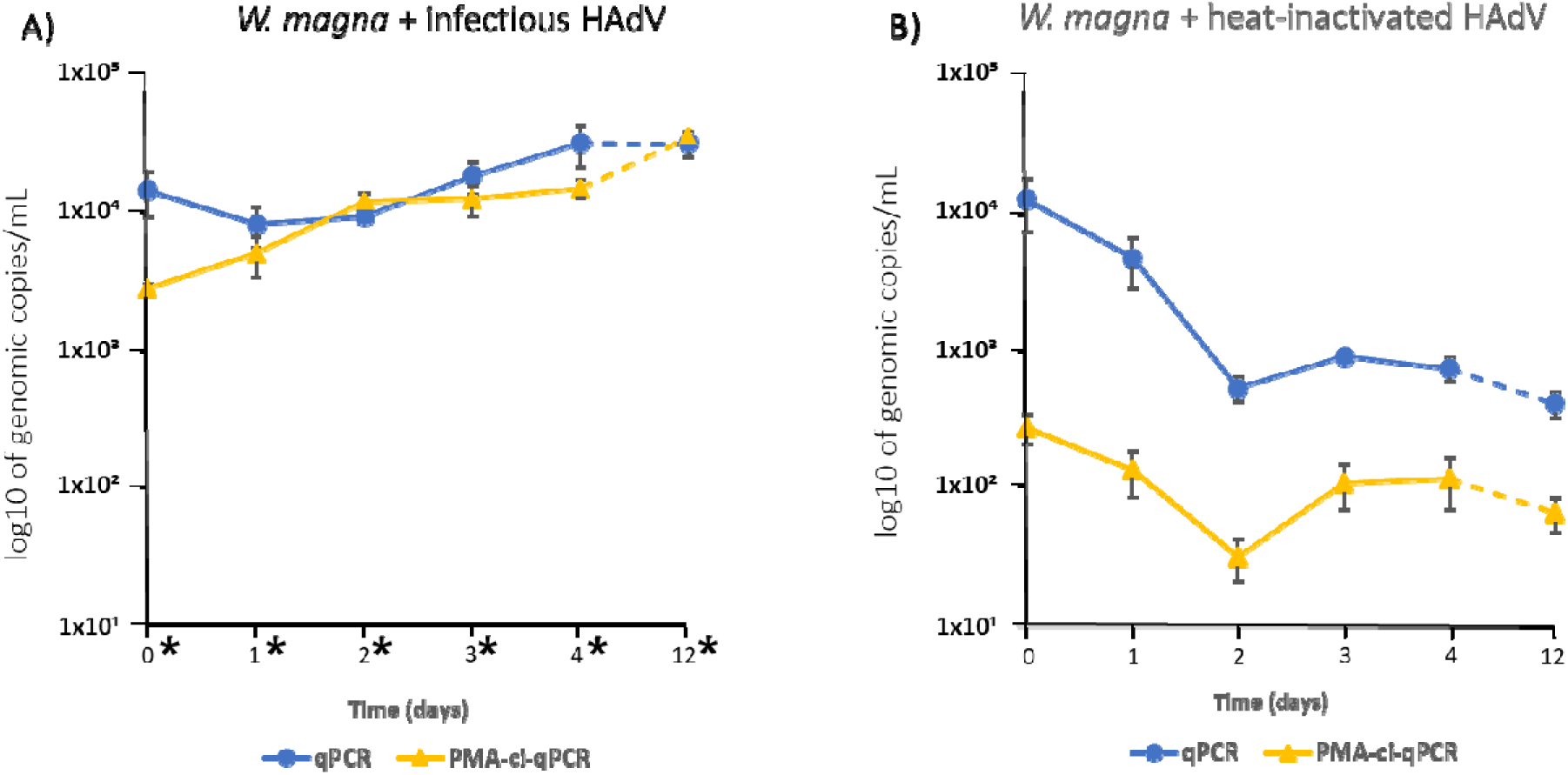
Molecular evaluation of HAdV replication activity using ICC-qPCR and PMA-ci-qPCR for infectious (A) and heat-inactivated (B) co-cultures. Unlike for heat-inactivated, inoculation of W. magna with infectious HAdV virions showed no reduction of genomic copies in both ICC-qPCR and PMA-ci-qPCR. After 12 days, a slight increase of viruses with an intact capsid can be detected after pre-treatment of the samples with the azo-dye PMA. Additionally, the presence of mRNA that is closely associated with virus replication and the formation of HAdV fibers could be detected above the limit of detection on all days. A loss of signal and total absence of mRNA was observed for heat inactivated HAdV, on the other hand. Error bars indicate the standard deviation, asterisk (*) the detection of HAdV mRNA (n=10).

## Discussion

The ability to avoid intracellular lysis, remain infectious or even replicate inside of FLA has been shown for numerous amoeba-resisting bacteria, such as with Legionella pneumophila as the most relevant opportunistic public health concern in engineered water system (Conza et al., 2013; Muchesa et al., 2017; Shaheen et al., 2019). Similar observations have been made for the giant amoebal viruses such as mimivirus (Arthofer et al., 2022; Scheid, 2015), and persist without amplification for various human enteric viruses, including reovirus (Dey et al., 2021a), coxsackievirus, adenovirus (Atanasova et al., 2018; Scheid and Schwarzenberger, 2012; Verani et al., 2016) and potentially rotavirus (Gad et al., 2019), along with respiratory viruses and their surrogates (Dey et al., 2021b; Folkins et al., 2020a). In avoiding digestion by their amoebae predators, viruses benefit from an additional layer of protection against environmental stressors and engineered disinfection processes. In turn, this has implications for their removal rates by sewage and drinking water treatment (Barrios-Hernández et al., 2020; Lodder et al., 2013; Qiu et al., 2015; Wang et al., 2018), as well as their ‘packaging’ and overall infectious dose at points of exposures.

While any one virus is considered to have a limited host range, they are generally expected to remain dormant, be destroyed or cause a cytopathic. On the other hand, viruses are the oldest parasites and may well have evolved with FLA for well over a billion years, providing a pandoras box of variants (Greub and Raoult, 2003; Oliveira et al., 2019a; Willemsen et al., 2025). Hence, we first explored potential cytopathic effects of HAdV 41/42 and HNoV GII.4 (the viral agents responsible for a large proportion in burden of disease due to food- and waterborne viral gastroenteritis (Carlson et al., 2024)). As illustrated in Figures 1 & 2 the three FLA species examined displayed neither cell damage nor changed growth rates in the presence of HAdV or HNoV. Examination using TEM and ImageStream™ flow cytometry coupled with fluorescent dyes further demonstrated clusters of characteristic virions in FLA organelles, vacuoles and within extracellular vesicles (Figures 3–7). Similar virion clustering has also been described by Atanasova et al. (2018) for coxsackievirus B5 in V. vermiformis and by Dey (2021b)– for respiratory viruses. In all cases, significant numbers of infectious virions were present in FLA trophozoites, cysts and excreted vesicles, likely increasing the actual dose delivered for any one human exposure event. Of added significant for public health, these packaged virions within cysts would result in extremely robust protection from natural and engineered disinfection systems (Folkins et al., 2020), extending viability for many months, only to release infectious virions once they excyst (Atanasova et al., 2018).

While the absence of a commercially available cell line for HNoV does not allow for the evaluation of viral infectivity after incorporation in V. vermiformis, A. polyphaga and W. magna, qPCR indicated that virus RNA concentrations in co-cultures with FLA remained stable for at least five days whereas free HNoV virons steadily (but slowly) decreased in the same time frame. As FLA actively graze on microorganisms and enzymatically digest them in food vacuoles, the apparent stability of HNoV dsRNA within FLA along with no change in FLA numbers, strongly implies it is not incorporated as a food source (Rodríguez-Zaragoza, 1994). Similar behaviour described by Hsueh and Gibson (2015) for the closely related murine norovirus (MNV) further implies the ability of caliciviruses to avoid digestion by FLA and maintain their infectivity (unlike for HNoV, the established RAW264.7 cell culture assay is available for MNV (Hwang et al., 2014).

The potential interaction between certain Acanthamoeba and human adenovirus has been investigated for more than a decade and the role of A. polyphaga in shielding HAdV from disinfection treatment has been shown by Verani et al. (2016) among others. Besides HAdV’s role in causing virus gastroenteritis in children in LMICs, they are also frequently mentioned as a potential sewage and technical performance indicator for rivers, lakes, beaches and engineered water systems (Farkas et al., 2020; Gerba and Betancourt, 2019; Rames et al., 2016; Saguti et al., 2022). A key attribute that makes HAdV a reliable indicator of sewage contamination is its presumed inability to replicate in aquatic environments. This characteristic reduces the risk of “false positive” signals arising from naturally occurring (autochthonous) microbial communities, thereby enhancing the accuracy of water quality assessments. Herein our work puts that assumption in question, as recently also observed for coliphages in river sediments, another promising group of faecal indicators and process surrogates (Mackowiak et al., 2018). Hence, care may be needed in interpreting detects, as it is for environmental E. coli that grow in the environment (Davies et al., 1995; Bertone et al., 2019).

Due to structural similarities between human macrophages and FLA like A. polyphaga, the hypothesis of possible replication of HAdV inside their amoeba host was evaluated but disproven by Scheid and Schwarzenberger (2012) and Maisonneuve et al. (2017). However, as described by Thomas et al. (2010), various members of the vast group of FLA can show significantly different behaviour when encountering fungi, bacteria and also viruses. Willaertia magna, closely related to Naegleria fowleri has moved into focus for amoeba-virus interaction due to increased examples of giant virus hosting (Ogata, 2018; Oliveira et al., 2019b) and maintenance of bacteriophages (Dey et al., 2021a; Dey et al., 2021b).

Co-culture with infectious and non-infectious HAdV virions revealed the entrance of virus particles not only into amoeba organelles but putatively into the amoeba nucleus (see Figure 5). Molecular assays targeting both the hexon gene region and mRNA associated with the fiber gene showed that culture containing infectious viruses maintained and increased genome copies over 12 days while qPCR for heat-inactivated HAdV reduced signal intensity (see Figure 8). The application of PMA based capsid integrity qPCR and introduction of the viruses into the integrated cell culture qPCR further indicated that HAdV not only avoided consumption by W. magna but maintained or even increased the number of infectious particles after 12 days. The presence of highly instable mRNA associated with the formation of novel virions as shown by Rodríguez et al. (2013) and Ko et al. (2005) throughout the co-culture, indicates some of kind activity resembling that of virus associated replication observable in their respective tissue culture host cells (Kuroshima et al., 2022).

Overall, these findings support the hypotheses that V. vermiformis, A. polyphaga, and W. magna internalize and allow for persistence of HNoV virions and the potential of HAdV to utilize the metabolism of W. magna to initiate virus replication. This and the microscopical visualization of HNoV and HAdV internalization and persistence inside of FLA potentially provide both enteric viruses a protective vessel to bypass widely used water disinfection treatments such as UV, chlorine or monochloramine, all of which have already been suspected to be insufficient to remove high virus loads (Barrios-Hernández et al., 2020; Leifels et al., 2016; Strathmann et al., 2016; Teixeira et al., 2020).

The lack of cell culture assays readily available to determine HNoV infectivity makes it difficult to evaluate the human health relevance of noroviruses incorporated and released by FLA. Capsid integrity assays could be used to indicate if the virus capsid is intact and the virion most likely capable of causing an infection. Ci-qPCR assays developed by Randazzo et al. (2018) are a useful tool to remove false-positive signals originating from inactivated viruses (Leifels et al., 2021). Application of this method to quantify HNoV with and without co-cultured FLA could allow for valuable insights into the effect of amoeba incorporation. If combined with established virus inactivation methods like hypochlorite, monochloramine, ozone or UV, it could further enhance our understanding about the implication of this virus-FLA cooperation in the context of wastewater and drinking water treatment.

## References

Anwar, A., Khan, N.A., Siddiqui, R., 2018. Combating Acanthamoeba spp. cysts: what are the options? Parasites & Vectors 11, 26.

Arthofer, P., Delafont, V., Willemsen, A., Panhölzl, F., Horn, M., 2022. Defensive symbiosis against giant viruses in amoebae. Proceedings of the National Academy of Sciences 119, e2205856119.

Ashbolt, N.J., 2023. Conceptual model to inform Legionella-amoebae control, including the roles of extracellular vesicles in engineered water system infections. Front Cell Infect Microbiol 13, 1200478.

Atanasova, N.D., Dey, R., Scott, C., Li, Q., Pang, X.-L., Ashbolt, N.J., 2018. Persistence of infectious Enterovirus within free-living amoebae–A novel waterborne risk pathway? Water research 144, 204–214.

Atmar, R.L., Opekun, A.R., Gilger, M.A., Estes, M.K., Crawford, S.E., Neill, F.H., Graham, D.Y., 2008. Norwalk virus shedding after experimental human infection. Emerg Infect Dis 14, 1553–1557.

Bányai, K., Estes, M.K., Martella, V., Parashar, U.D., 2018. Viral gastroenteritis. The Lancet 392, 175–186.

Barrios-Hernández, M.L., Pronk, M., Garcia, H., Boersma, A., Brdjanovic, D., van Loosdrecht, M.C.M., Hooijmans, C.M., 2020. Removal of bacterial and viral indicator organisms in full-scale aerobic granular sludge and conventional activated sludge systems. Water Res X 6, 100040.

Buse, H.Y., Ji, P., Gomez-Alvarez, V., Pruden, A., Edwards, M.A., Ashbolt, N.J., 2017. Effect of temperature and colonization of Legionella pneumophila and Vermamoeba vermiformis on bacterial community composition of copper drinking water biofilms. Microb Biotechnol 10, 773–788.

Carlson, K.B., Dilley, A., O’Grady, T., Johnson, J.A., Lopman, B., Viscidi, E., 2024. A narrative review of norovirus epidemiology, biology, and challenges to vaccine development. npj Vaccines 9, 94.

Cassini, A., Colzani, E., Kramarz, P., Kretzschmar, M.E., Takkinen, J., 2016. Impact of food and water-borne diseases on European population health. Current Opinion in Food Science 12, 21–29.

Chaúque, B.J.M., Rott, M.B., 2022. The role of free-living amoebae in the persistence of viruses in the era of severe acute respiratory syndrome 2, should we be concerned? Rev Soc Bras Med Trop 55, e0045.

Chen, L., Jiao, J., Liu, S., Liu, L., Liu, P., 2023. Mapping the global, regional, and national burden of diarrheal diseases attributable to unsafe water. Front Public Health 11, 1302748.

Collier, S.A., Deng, L., Adam, E.A., Benedict, K.M., Beshearse, E.M., Blackstock, A.J., Bruce, B.B., Derado, G., Edens, C., Fullerton, K.E., Gargano, J.W., Geissler, A.L., Hall, A.J., Havelaar, A.H., Hill, V.R., Hoekstra, R.M., Reddy, S.C., Scallan, E., Stokes, E.K., Yoder, J.S., Beach, M.J., 2021. Estimate of Burden and Direct Healthcare Cost of Infectious Waterborne Disease in the United States. Emerg Infect Dis 27, 140–149.

Colson, P., Aherfi, S., La Scola, B., 2017. Evidence of giant viruses of amoebae in the human gut. Human Microbiome Journal 5–6, 14-19.

Conza, L., Pagani, S.C., Gaia, V., 2013. Presence of Legionella and Free-Living Amoebae in Composts and Bioaerosols from Composting Facilities. PLOS ONE 8, e68244.

da Silva, T.C.B., Chaúque, B.J.M., Benitez, G.B., Rott, M.B., 2024. Global prevalence of potentially pathogenic free-living amoebae in sewage and sewage-related environments—systematic review with meta-analysis. Parasitol Res 123, 148.

Dey, R., Dlusskaya, E., Ashbolt, N.J., 2021a. SARS-CoV-2 surrogate (Phi6) environmental persistence within free-living amoebae. Journal of Water and Health 20, 83–91.

Dey, R., Folkins, M.A., Ashbolt, N.J., 2021b. Extracellular amoebal-vesicles: potential transmission vehicles for respiratory viruses. npj Biofilms and Microbiomes 7, 25.

Fan, S., Shen, Y., Qian, L., 2024. Social life of free-living amoebae in aquatic environment— comprehensive insights into interactions of free-living amoebae with neighboring microorganisms. Frontiers in Microbiology 15.

Farkas, K., Mannion, F., Hillary, L.S., Malham, S.K., Walker, D.I., 2020. Emerging technologies for the rapid detection of enteric viruses in the aquatic environment. Current Opinion in Environmental Science & Health 16, 1–6.

Flemming, H.C., Wingender, J., 2010. The biofilm matrix. Nat Rev Microbiol 8, 623–633.

Flemming, H.C., Wuertz, S., 2019. Bacteria and archaea on Earth and their abundance in biofilms. Nature Reviews Microbiology 17, 247–260.

Folkins, M.A., Dey, R., Ashbolt, N.J., 2020a. Interactions between Human Reovirus and Free-Living Amoebae: Implications for Enteric Virus Disinfection and Aquatic Persistence. Environmental Science & Technology 54, 10201–10206.

Folkins, M.A., Dey, R., Ashbolt, N.J., 2020b. Interactions between Human Reovirus and Free-Living Amoebae: Implications for Enteric Virus Disinfection and Aquatic Persistence. Environ Sci Technol 54, 10201–10206.

Free, R.J., Buss, B.F., Koirala, S., Ulses, M., Carlson, A., Loeck, B., Safranek, T., 2019. Successive Norovirus Outbreaks at an Event Center - Nebraska, October-November, 2017. MMWR. Morbidity and mortality weekly report 68, 627–630.

Gad, M., Allayeh, A., Elmahdy, E., Shaheen, M., Rizk, N., Al-Herrawy, A., Saleh Ibrahim, F.E.-Z., Saleh, R., Ali, M., 2019. Genotyping and interaction-reality of Acanthamoeba, enteric adenovirus and rotavirus in drinking water, Egypt. ARTICLE INFO ABSTRACT. Egyptian Journal of Aquatic Biology and Fisheries 32, 65–79.

Gerba, C.P., Betancourt, W.Q., 2019. Assessing the Occurrence of Waterborne Viruses in Reuse Systems: Analytical Limits and Needs. Pathogens (Basel, Switzerland) 8.

Girardi, V., Demoliner, M., Gularte, J.S., Spilki, F.R., 2019. ’Don’t put your head under water’: enteric viruses in Brazilian recreational waters. New Microbes New Infect 29, 100519–100519.

Gómez-Gómez, C., Ramos-Barbero, M.D., Sala-Comorera, L., Morales-Cortes, S., Vique, G., García-Aljaro, C., Muniesa, M., 2024. Persistence of crAssBcn phages in conditions of natural inactivation and disinfection process and their potential role as human source tracking markers. Science of The Total Environment 957, 177450.

Greub, G., Raoult, D., 2003. History of the ADP/ATP-translocase-encoding gene, a parasitism gene transferred from a Chlamydiales ancestor to plants 1 billion years ago. Appl Environ Microbiol 69, 5530–5535.

Hall, A.J., 2012. Noroviruses: the perfect human pathogens? J Infect Dis 205, 1622–1624.

Hamilton, K.A., Ahmed, W., Palmer, A., Sidhu, J.P.S., Hodgers, L., Toze, S., Haas, C.N., 2016. Public health implications of Acanthamoeba and multiple potential opportunistic pathogens in roof-harvested rainwater tanks. Environ Res 150, 320–327.

Heim, A., Ebnet, C., Harste, G., Pring Åkerblom, P., 2003. Rapid and quantitative detection of human adenovirus DNA by real time PCR. Journal of medical virology 70, 228–239.

Hsueh, T.-Y., Gibson, K.E., 2015. Interactions between Human Norovirus Surrogates and <span class=“named-content genus-species” id=“named-content-1“>Acanthamoeba</span> spp. Applied and Environmental Microbiology 81, 4005–4013.

Hutson, A.M., Atmar, R.L., Estes, M.K., 2004. Norovirus disease: changing epidemiology and host susceptibility factors. Trends Microbiol 12, 279–287.

Hwang, S., Alhatlani, B., Arias, A., Caddy, S.L., Christodoulou, C., Cunha, J.B., Emmott, E., Gonzalez-Hernandez, M., Kolawole, A., Lu, J., Rippinger, C., Sorgeloos, F., Thorne, L., Vashist, S., Goodfellow, I., Wobus, C.E., 2014. Murine norovirus: propagation, quantification, and genetic manipulation. Curr Protoc Microbiol 33, 15K.12.11-15K.12.61.

ISO 2017. ISO 15216-21. Microbiology of the food chain – Horizontal method for determination of hepatitis A virus and norovirus using real-time RT-PCR. .

Kageyama, T., Kojima, S., Shinohara, M., Uchida, K., Fukushi, S., Hoshino, F.B., Takeda, N., Katayama, K., 2003. Broadly reactive and highly sensitive assay for Norwalk-like viruses based on real-time quantitative reverse transcription-PCR. Journal of clinical microbiology 41, 1548–1557.

Kebbi-Beghdadi, C., Greub, G., 2014. Importance of amoebae as a tool to isolate amoeba-resisting microorganisms and for their ecology and evolution: the Chlamydia paradigm. Environ Microbiol Rep 6, 309–324.

Khales, P., Razizadeh, M.H., Ghorbani, S., Moattari, A., Sarvari, J., Saadati, H., Sayyahfar, S., Salavatiha, Z., Hasanabad, M.H., Poortahmasebi, V., Tavakoli, A., 2024. Human adenoviruses in children with gastroenteritis: a systematic review and meta-analysis. BMC Infectious Diseases 24, 478.

Ko, G., Jothikumar, N., Hill, V.R., Sobsey, M.D., 2005. Rapid detection of infectious adenoviruses by mRNA real-time RT-PCR. Journal of virological methods 127, 148–153.

Kuroshima, T., Matsuda, A.Y., Hossain, E., Yasuda, M., Kitamura, T., Kitagawa, Y., Higashino, F., 2022. Adenovirus infection controls processing bodies to stabilize AU-rich element-containing mRNA. Virology 573, 124–130.

Lee, N., Chan, M.C.W., Wong, B., Choi, K.W., Sin, W., Lui, G., Chan, P.K.S., Lai, R.W.M., Cockram, C.S., Sung, J.J.Y., Leung, W.K., 2007. Fecal viral concentration and diarrhea in norovirus gastroenteritis. Emerging infectious diseases 13, 1399–1401.

Lei, H., Li, Y., Xiao, S., Yang, X., Lin, C., Norris, S.L., Wei, D., Hu, Z., Ji, S., 2017. Logistic growth of a surface contamination network and its role in disease spread. Scientific Reports 7, 14826.

Leifels, M., Cheng, D., Sozzi, E., Shoults, D.C., Wuertz, S., Mongkolsuk, S., Sirikanchana, K., 2021. Capsid integrity quantitative PCR to determine virus infectivity in environmental and food applications - A systematic review. Water Res X 11, 100080.

Leifels, M., Hamza, I.A., Krieger, M., Wilhelm, M., Mackowiak, M., Jurzik, L., 2016. From Lab to Lake - Evaluation of Current Molecular Methods for the Detection of Infectious Enteric Viruses in Complex Water Matrices in an Urban Area. PLoS One 11, e0167105.

Leifels, M., Shoults, D., Wiedemeyer, A., Ashbolt, N.J., Sozzi, E., Hagemeier, A., Jurzik, L., 2019. Capsid Integrity qPCR—An Azo-Dye Based and Culture-Independent Approach to Estimate Adenovirus Infectivity after Disinfection and in the Aquatic Environment. Water 11, 1196.

Lodder, W.J., Rutjes, S.A., Takumi, K., de Roda Husman, A.M., 2013. Aichi virus in sewage and surface water, the Netherlands. Emerging infectious diseases 19, 1222–1230.

Loisy, F., Atmar, R.L., Guillon, P., Le Cann, P., Pommepuy, M., Le Guyader, F.S., 2005. Real-time RT-PCR for norovirus screening in shellfish. Journal of virological methods 123, 1–7.

Mackowiak, M., Leifels, M., Hamza, I.A., Jurzik, L., Wingender, J., 2018. Distribution of Escherichia coli, coliphages and enteric viruses in water, epilithic biofilms and sediments of an urban river in Germany. Science of the Total Environment 626, 650–659.

Maisonneuve, E., Cateau, E., Leveque, N., Kaaki, S., Beby-Defaux, A., Rodier, M.-H., 2017. Acanthamoeba castellanii is not be an adequate model to study human adenovirus interactions with macrophagic cells. PloS one 12, e0178629–e0178629.

Mattana, A., Serra, C., Mariotti, E., Delogu, G., Fiori, P.L., Cappuccinelli, P., 2006. Acanthamoeba castellanii promotion of in vitro survival and transmission of coxsackie b3 viruses. Eukaryotic Cell 5, 665–671.

Muchesa, P., Leifels, M., Jurzik, L., Barnard, T.G., Bartie, C., 2016. Free-living amoebae isolated from a hospital water system in South Africa: a potential source of nosocomial and occupational infection. Water Sci Tech-W Sup 16, 70–78.

Muchesa, P., Leifels, M., Jurzik, L., Hoorzook, K.B., Barnard, T.G., Bartie, C., 2017. Coexistence of free-living amoebae and bacteria in selected South African hospital water distribution systems. Parasitol Res 116, 155–165.

O’Shea, H., Blacklaws, B.A., Collins, P.J., McKillen, J., Fitzgerald, R., 2019. Viruses Associated With Foodborne Infections. Reference Module in Life Sciences, B978-970-912-809633-809638.890273-809635.

Ogata, H., 2018. Habitat Alterations by Viruses: Strategies by Tupanviruses and Others. Microbes Environ 33, 117–119.

Oliveira, G., La Scola, B., Abrahão, J., 2019a. Giant virus vs amoeba: fight for supremacy. Virology Journal 16, 126.

Oliveira, G., Silva, L., Leão, T., Mougari, S., da Fonseca, F.G., Kroon, E.G., La Scola, B., Abrahão, J.S., 2019b. Tupanvirus-infected amoebas are induced to aggregate with uninfected cells promoting viral dissemination. Scientific Reports 9, 183.

Qiu, Y., Lee, B.E., Neumann, N., Ashbolt, N., Craik, S., Maal-Bared, R., Pang, X.L., 2015. Assessment of human virus removal during municipal wastewater treatment in Edmonton, Canada. Journal of Applied Microbiology 119, 1729–1739.

Rames, E., Roiko, A., Stratton, H., Macdonald, J., 2016. Technical aspects of using human adenovirus as a viral water quality indicator. Water Research 96, 308–326.

Randazzo, W., Khezri, M., Ollivier, J., Le Guyader, F.S., Rodriguez-Diaz, J., Aznar, R., Sanchez, G., 2018. Optimization of PMAxx pretreatment to distinguish between human norovirus with intact and altered capsids in shellfish and sewage samples. Int J Food Microbiol 266, 1–7.

Rodríguez-Zaragoza, S., 1994. Ecology of Free-Living Amoebae. Critical Reviews in Microbiology 20, 225–241.

Rodríguez, R.A., Polston, P.M., Wu, M.J., Wu, J., Sobsey, M.D., 2013. An improved infectivity assay combining cell culture with real-time PCR for rapid quantification of human adenoviruses 41 and semi-quantification of human adenovirus in sewage. Water Research 47, 3183–3191.

Saguti, F., Churqui, M.P., Kjellberg, I., Wang, H., Ottoson, J., Paul, C., Bergstedt, O., Norder, H., Nyström, K., 2022. The UV Dose Used for Disinfection of Drinking Water in Sweden Inadequately Inactivates Enteric Virus with Double-Stranded Genomes. Int J Environ Res Public Health 19.

Samba-Louaka, A., Delafont, V., Rodier, M.-H., Cateau, E., Héchard, Y., 2019. Free-living amoebae and squatters in the wild: ecological and molecular features. FEMS Microbiology Reviews 43, 415–434.

Scallan Walter, E., Cui, Z., Tierney, R., Griffin, P., Hoekstra, R., Payne, D., Rose, E., Devine, C., Namwase, A.S., Mirza, S., Kambhampati, A., Straily, A., Bruce, B., 2025. Foodborne Illness Acquired in the United States—Major Pathogens, 2019. Emerging Infectious Disease journal 31, 669.

Scheid, P., 2015. Viruses in close associations with free-living amoebae. Parasitol Res 114, 3959–3967.

Scheid, P., Schwarzenberger, R., 2012. Acanthamoeba spp. as vehicle and reservoir of adenoviruses. Parasitol Res 111, 479–485.

Shaheen, M., Ashbolt, N.J., 2018. Free-Living Amoebae Supporting Intracellular Growth May Produce Vesicle-Bound Respirable Doses of Legionella Within Drinking Water Systems. Exposure and Health 10, 201–209.

Shaheen, M., Scott, C., Ashbolt, N.J., 2019. Long-term persistence of infectious Legionella with free-living amoebae in drinking water biofilms. International journal of hygiene and environmental health 222, 678–686.

Shmakova, L., Bondarenko, N., Smirnov, A., 2016. Viable Species of Flamella (Amoebozoa: Variosea) Isolated from Ancient Arctic Permafrost Sediments. Protist 167, 13–30.

Storey, M.V., Winiecka-Krusnell, J., Ashbolt, N.J., Stenström, T.A., 2004. The efficacy of heat and chlorine treatment against thermotolerant Acanthamoebae and Legionellae. Scand J Infect Dis 36, 656–662.

Strathmann, M., Horstkott, M., Koch, C., Gayer, U., Wingender, J., 2016. The River Ruhr - an urban river under particular interest for recreational use and as a raw water source for drinking water: The collaborative research project “Safe Ruhr” - microbiological aspects. International journal of hygiene and environmental health 219, 643–661.

Sun, S., Noorian, P., McDougald, D., 2018. Dual Role of Mechanisms Involved in Resistance to Predation by Protozoa and Virulence to Humans. Frontiers in Microbiology 9.

Teixeira, P., Costa, S., Brown, B., Silva, S., Rodrigues, R., Valério, E., 2020. Quantitative PCR Detection of Enteric Viruses in Wastewater and Environmental Water Sources by the Lisbon Municipality: A Case Study. Water 12, 544.

Thomas, J.M., Ashbolt, N.J., 2011. Do Free-Living Amoebae in Treated Drinking Water Systems Present an Emerging Health Risk? Environmental Science & Technology 45, 860–869.

Thomas, V., McDonnell, G., Denyer, S.P., Maillard, J.-Y., 2010. Free-living amoebae and their intracellular pathogenic microorganisms: risks for water quality. FEMS microbiology reviews 34, 231–259.

Troeger, C., Blacker, B.F., Khalil, I.A., Rao, P.C., Cao, S., Zimsen, S.R.M., Albertson, S.B., Stanaway, J.D., Deshpande, A., Abebe, Z., Alvis-Guzman, N., Amare, A.T., Asgedom, S.W., Anteneh, Z.A., Antonio, C.A.T., Aremu, O., Asfaw, E.T., Atey, T.M., Atique, S., Avokpaho, E.F.G.A., Awasthi, A., Ayele, H.T., Barac, A., Barreto, M.L., Bassat, Q., Belay, S.A., Bensenor, I.M., Bhutta, Z.A., Bijani, A., Bizuneh, H., Castañeda-Orjuela, C.A., Dadi, A.F., Dandona, L., Dandona, R., Do, H.P., Dubey, M., Dubljanin, E., Edessa, D., Endries, A.Y., Eshrati, B., Farag, T., Feyissa, G.T., Foreman, K.J., Forouzanfar, M.H., Fullman, N., Gething, P.W., Gishu, M.D., Godwin, W.W., Gugnani, H.C., Gupta, R., Hailu, G.B., Hassen, H.Y., Hibstu, D.T., Ilesanmi, O.S., Jonas, J.B., Kahsay, A., Kang, G., Kasaeian, A., Khader, Y.S., Khalil, I.A., Khan, E.A., Khan, M.A., Khang, Y.-H., Kissoon, N., Kochhar, S., Kotloff, K.L., Koyanagi, A., Kumar, G.A., Magdy Abd El Razek, H., Malekzadeh, R., Malta, D.C., Mehata, S., Mendoza, W., Mengistu, D.T., Menota, B.G., Mezgebe, H.B., Mlashu, F.W., Murthy, S., Naik, G.A., Nguyen, C.T., Nguyen, T.H., Ningrum, D.N.A., Ogbo, F.A., Olagunju, A.T., Paudel, D., Platts-Mills, J.A., Qorbani, M., Rafay, A., Rai, R.K., Rana, S.M., Ranabhat, C.L., Rasella, D., Ray, S.E., Reis, C., Renzaho, A.M.N., Rezai, M.S., Ruhago, G.M., Safiri, S., Salomon, J.A., Sanabria, J.R., Sartorius, B., Sawhney, M., Sepanlou, S.G., Shigematsu, M., Sisay, M., Somayaji, R., Sreeramareddy, C.T., Sykes, B.L., Taffere, G.R., Topor-Madry, R., Tran, B.X., Tuem, K.B., Ukwaja, K.N., Vollset, S.E., Walson, J.L., Weaver, M.R., Weldegwergs, K.G., Werdecker, A., Workicho, A., Yenesew, M., Yirsaw, B.D., Yonemoto, N., El Sayed Zaki, M., Vos, T., Lim, S.S., Naghavi, M., Murray, C.J.L., Mokdad, A.H., Hay, S.I., Reiner, R.C., Jr., 2018. Estimates of the global, regional, and national morbidity, mortality, and aetiologies of diarrhoea in 195 countries: a systematic analysis for the Global Burden of Disease Study 2016. The Lancet Infectious Diseases 18, 1211–1228.

Verani, M., Di Giuseppe, G., Tammaro, C., Carducci, A., 2016. Investigating the role of Acanthamoeba polyphaga in protecting Human Adenovirus from water disinfection treatment. European Journal of Protistology 54, 11–18.

Walker, D.I., Witt, J., Rostant, W., Burton, R., Davison, V., Ditchburn, J., Evens, N., Godwin, R., Heywood, J., Lowther, J.A., Peters, N., Porter, J., Posen, P., Wickens, T., Wade, M.J., 2024. Piloting wastewater-based surveillance of norovirus in England. Water Res 263, 122152.

Wang, H., Sikora, P., Rutgersson, C., Lindh, M., Brodin, T., Björlenius, B., Larsson, D.G.J., Norder, H., 2018. Differential removal of human pathogenic viruses from sewage by conventional and ozone treatments. International journal of hygiene and environmental health 221, 479–488.

Willemsen, A., Manzano-Marín, A., Horn, M., 2025. Novel High-Quality Amoeba Genomes Reveal Widespread Codon Usage Mismatch Between Giant Viruses and Their Hosts. Genome Biol Evol 17.

Zhang, M., Altan-Bonnet, N., Shen, Y., Shuai, D., 2022. Waterborne Human Pathogenic Viruses in Complex Microbial Communities: Environmental Implication on Virus Infectivity, Persistence, and Disinfection. Environmental Science and Technology 56, 5381–5389.

Ziros, P.G., Kokkinos, P.A., Allard, A., Vantarakis, A., 2015. Development and Evaluation of a Loop-Mediated Isothermal Amplification Assay for the Detection of Adenovirus 40 and 41. Food and environmental virology 7, 276–285.

